# Genetic and microbial analysis of invasiveness for *Escherichia coli* strains associated with inflammatory bowel disease

**DOI:** 10.1101/2024.09.23.613251

**Authors:** Jungyeon Kim, Jing Zhang, Lisa Kinch, Jinhui Shen, Sydney Field, Jan-Michael Klapproth, Kevin J. Forsberg, Tamia Harris-Tryon, Kim Orth, Qian Cong, Josephine Ni

## Abstract

**Background & Aims:** The adherent-invasive *Escherichia coli* (AIEC) pathotype is implicated in inflammatory bowel disease (IBD) pathogenesis. AIEC strains are currently defined by phenotypic measurement of their pathogenicity, including invasion of epithelial cells. This broad definition, combined with the genetic diversity of AIEC across IBD patients, has complicated the identification of virulence determinants. We sought to quantify the invasion phenotype of clinical isolates from IBD patients and identify the genetic basis for their invasion into epithelial cells.

**Methods:** A pangenome with core and accessory genes (genotype) was assembled using whole genome sequencing of 168 *E. coli* samples isolated from 13 IBD patients. A modified assay for invasion of epithelial cells (phenotype) was established with consideration of antibiotic resistance phenotypes. Isolate genotype was correlated to invasiveness phenotype to identify genetic factors that co-segregate with invasion.

**Results:** Pangenome-wide comparisons of *E. coli* clinical isolates identified accessory genes that can co-segregate with invasion phenotype. These correlations found the acquisition of antibiotic resistance genes in clinical isolates compromised the traditional gentamicin protection assays used to quantify invasion. Therefore, an alternate assay, based on amikacin resistance, identified genes co-segregating with invasion. These genes encode an arylsulfatase, a glycoside hydrolase, and genetic islands carrying propanediol utilization and sulfoquinovose metabolism pathways.

**Conclusions:** This study highlights the importance of incorporating antibiotic resistance screening for invasion assays used in AIEC identification. Accurately screened invasion phenotypes identified accessory genome elements among *E. coli* IBD isolates that correlate with their ability to invade epithelial cells. These results help explain why single genetic markers for the AIEC phylotype are challenging to identify.

**Synopsis:** We established a pangenome of *Escherichia coli* isolates from inflammatory bowel disease patients using whole genome sequencing. The genotypes were correlated with a newly developed measurement of invasion phenotype to identify co-segregating genes.

## Introduction

*Escherichia coli* (*E. coli*) are common colonizers of the vertebrate digestive tract and appear frequently as commensal constituents of the human gut microbiome (1, 2). Many *E. coli* variants possess virulence factors that facilitate gut or urinary tract infections, even in otherwise healthy individuals (3, 4). The presence of these virulence factors and the replicative niche the bacteria colonize during infection forms the basis of the pathotype categorization of these *E. coli* (3). While the *E. coli* core genome, which determines the bacteria’s phylogenetic classification, is highly conserved across both pathogenic and commensal strains, pangenomic analyses reveal diversity within a highly variable accessory genome that distinguishes pathogenic *E. coli* both from their commensal counterparts and from one another (3, 5–7). Analyses of the accessory genome have likewise illuminated how horizontal genetic acquisition, genetic diversification, and gene loss have blurred the lines differentiating pathotypes (3, 6).

The AIEC pathotype has classically been characterized by bacterial invasion of the host intestinal epithelium and propagation within macrophages without inducing cell death (8–10). Although AIEC can also be identified in healthy individuals, they are consistently enriched in the microbiota of patients with inflammatory bowel disease (IBD), especially Crohn’s Disease (CD) (8, 10). They correlate with elevated gut inflammation and dysbiosis of the intestinal microbiome (11, 12). Further, they cause disease when inoculated into murine colitis models (13). Many efforts have been made to identify AIEC strains genotypically, but they exhibit significant diversity across individual CD patients (14). This diversity across clinical isolates presents a challenge in linking the AIEC pathotype with specific genetic markers. Few virulence factors have been identified in AIEC accessory genomes, and no consensus exists for genetic markers within the accessory genome (15, 16). Due to this limitation, the gold standard for identifying AIEC is a phenotypic gentamicin protection assay where intracellular bacterial invasion is quantified (17, 18). Unfortunately, *E. coli* isolates from CD patients exhibit elevated frequencies of antibiotic resistance relative to those from healthy individuals, partially because CD patients are immunocompromised and are frequently exposed to antibiotics as part of their treatment (19, 20).

Here, using 168 *E. coli* samples isolated from 13 IBD patients, we aimed to evaluate their invasiveness and correlate such ability with genetic markers. However, we found that gentamicin protection assays were not trustworthy as we observed frequent gains of gentamicin resistance genes, resulting in false positives for invasion. We thus developed an alternative amikacin protection assay that proved more reliable. Based on these data, we recommend 1) including an antibiotic resistance screen in the phenotypic identification of AIEC and 2) using the amikacin protection assays to complement the classic gentamicin protection assays, which may eliminate false positives and provide a more stringent filter for pathotype constituents. Integrating results from our improved assays and comparative genomic analyses, we identified genes that co-segregate with the invasiveness. These include genomic neighborhoods encoding propanediol utilization and sulfoquinovose (SQ) metabolism, as well as an arylsulfatase (AslA) and a glycoside hydrolase (GH127).

## Results

### Clinical *E. coli* Isolate collection and genomic sequencing

A total of 49 subjects undergoing colonoscopy consented to sample collection, and their demographics are summarized in **Table 1**. Bacterial growth on MacConkey agar plates was detected in the form of individual lactose-fermenting and nonfermenting colonies. 830 bacterial isolates were further analyzed in triplicate by an *in vitro* gentamicin protection assay with differentiated Caco-2 epithelial cells, including 378 isolates from CD patients, 111 from UC patients, and 341 from normal control subjects. From these assays, 117 AIEC were tentatively identified based on significant epithelial invasion defined as a percent invasion of >1% (21). Consistent with previous studies, more AIEC strains were found in CD patients (26%) versus UC patients (18%) and normal controls (13%) (21).

**Table 1.** Subject demographics.

|  | CD | UC | Indeterminate<br>Colitis | Controls |
| --- | --- | --- | --- | --- |
| Number of subjects | 23 | 12 | 2 | 12 |
| Female sex | 60% | 75% | 50% | 57% |
| Mean age | 44 | 38 | 54 | 60 |
| Years since diagnosis |  |  |  |  |
| Previous intestinal resection |  |  |  |  |
| 5-ASA treatment | 48% | 83% | 0% | - |
| 6MP/AZA treatment | 30% | 25% | 0% | - |
| Steroid treatment | 26% | 8% | 0% | - |
| Anti-TNF treatment | 17% | 17% | 0% | - |
| Antibiotics | 17% | 0% | 0% | - |

We attempted to obtain whole genome sequences using both Nanopore and Illumina platforms for 213 isolates from this clinical collection that were characterized as *E. coli* via biochemical testing with the Api-20E system. The Illumina short reads and Nanopore long reads were combined to assemble the genomes of these candidate *E. coli* isolates. Analysis of these assembled genomes by NCBI Prokaryotic Genome Annotation Pipeline revealed that 45 (21%) of them were not *E. coli* based on the genomic data, and instead, they were other microbes associated with humans, such as *Citrobacter rodentium* and *Klebsiella pneumoniae*.

Therefore, we focused on the remaining 168 *E. coli* isolates in our study. On average, we obtained 350 Mbp and 420 Mbp Illumina and Nanopore reads, respectively, for each isolate (**Table S1**).

These reads allowed us to assemble high-quality genomes for nearly all isolates: 154 (92%) were each assembled into one long scaffold corresponding to the chromosome and several short scaffolds corresponding to plasmids or regions difficult to assemble into the chromosome (**Table S1**). The assembled genomes for these isolates range between 4.7Mbp and 5.5Mbp, and they encode 4.3k to 5.1k proteins, typical for *E. coli*. We further assessed the quality of these assembled genomes by CheckM (22), which evaluates the completeness by the presence of a set of universal single-copy genes (USCGs) expected to be present in a lineage. CheckM also detects potential contaminations in genome assemblies by the presence of more than one copy of such USCGs. Our genome assemblies show a median completeness of 99.2% and contamination of 0.37%; both are better than the median values (99.0% and 0.59%) of high-quality bacterial genomes in NCBI.

### Clonal Nature of Patient Isolates

Phylogenetic analysis of all sequenced isolates highlighted their clonal nature among patients (**Fig 1A**). Similar to previous reports for *E. coli* isolates from CD patients (14, 20, 23), the isolates collected from different patients with CD, UC, and indeterminate colitis are phylogenetically diverse and span 6 phylogroups (A, B1, B2, D, E, and G; **Fig 1A**). However, isolates collected from the same patient were mostly clonal, regardless of the biopsy location. The clonal nature of isolates from the same patient is also reflected by the average sequence identity between orthologous segments (**Fig 1B**). While isolates from different patients show sequence identity between 96.3% and 99.1% (green dots in **Fig 1B**), pairs from the same patient mostly show 100% sequence identity (orange dots in **Fig 1B**).

**Figure 1.**
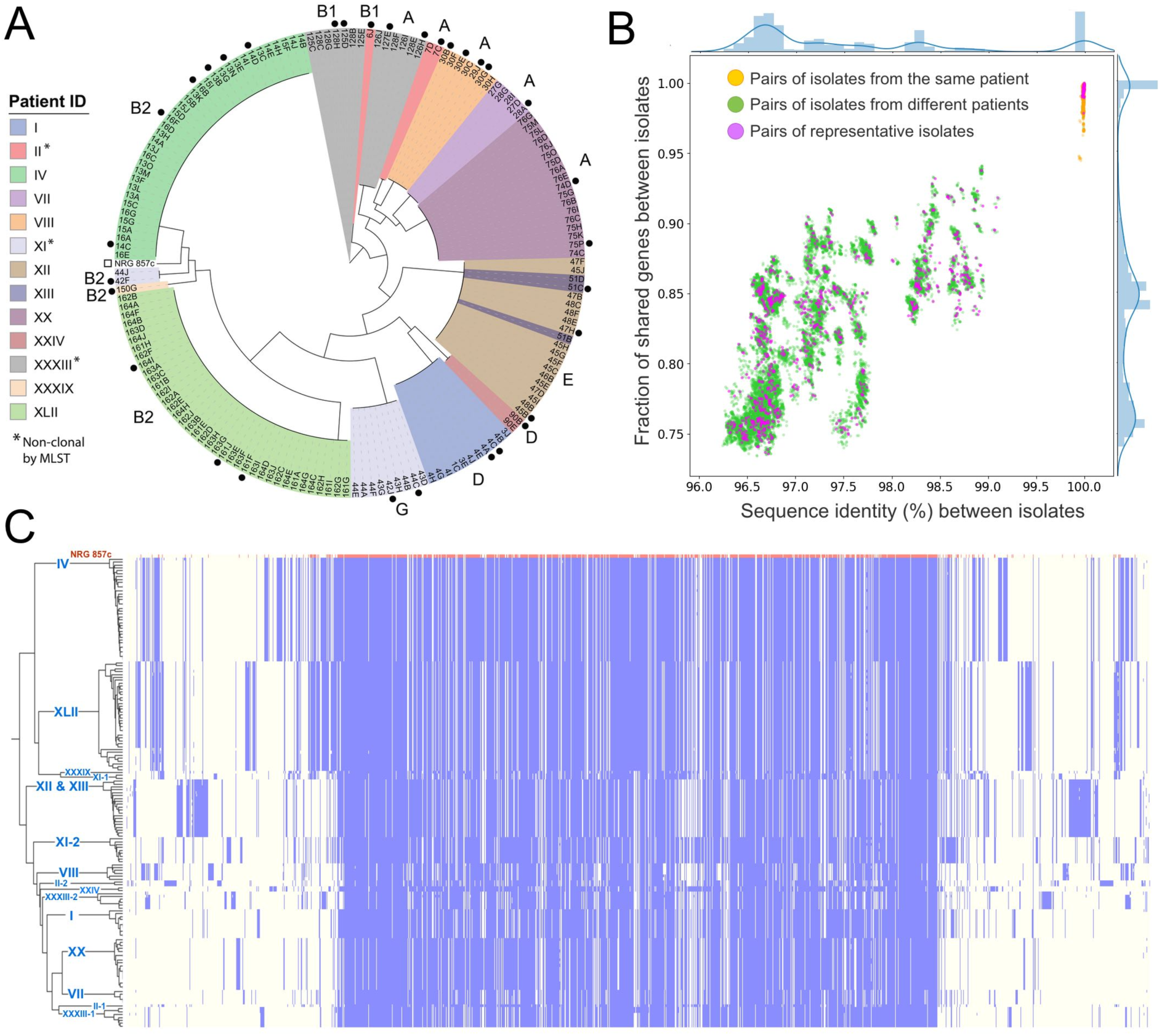
Pangenome of clonal patient *E. coli* isolates. **(A)** Maximum-likelihood phylogenetic tree based on the concatenated protein multiple sequence alignment of orthologous groups. Isolates are colored by patient id, and patients with non-clonal isolates by MLST typing are indicated by*. Selected reference isolates (black circle) and the public AIEC strain (open square) are indicated outside the tree. Phylotypes for each clonal group are labeled outside the tree. **(B)** The similarity between *E. coli* isolates is measured by the average nucleotide identity (X-axis) and the fraction of shared genes (Y-axis). Each dot in the scatterplot represents a pair of isolates, and the bar plots by the scatterplots show the distribution of values on these axes. **(C)** A heatmap showing the presence (in blue) and absence (in light yellow) of genes (X-axis) across different *E. coli* isolates (Y-axis). The genes are ordered along the axes based on their average genomic distances in all the isolates shown in the graph. The first row of the heatmap represents the reference strain, NRG 857c, while other rows show the *E. coli* isolates sequenced in this study. These isolates are ordered based on their phylogeny on the left of the heatmap. Patient IDs associated with these isolated are labeled on the tree branch.

The shared gene content among isolate genomes positively correlates with the sequence identity between isolates (**Fig 1B**). However, compared to sequence identity, the gene content of these isolates shows a much larger variability: isolates from different patients share 74% to 94% of genes, and isolates showing 100% sequence identity in orthologs may still possess slightly different sets (< 5%) of genes (**Fig 1B**). Such variability is common among bacterial isolates (24) and highlights the important role of horizontal gene transfer (HGT) in the evolution of microbes. As a result of HGT, the sequenced isolates process different sets of accessory genes in addition to the shared core genomes.

To characterize the diverse set of proteins encoded by the sequenced *E. coli* isolates, we identified orthologous groups (OGs) among these proteomes and assembled a pangenome reference protein set that includes one representative protein per OG (**Table S2**). The presence or absence of each OG in each isolate is shown as a heatmap in **Fig 1C**. In this heatmap, the reference proteins are ordered to approximate their genomic distances, placing each gene near its genomic neighbors. The core genes shared by all isolates are mostly encoded by the chromosome, and they cluster in the middle of this heatmap. In contrast, the accessory genes, frequently encoded on plasmids, are placed around the core genome in the heatmap. This distribution underscores that although core genes are shared by all isolates, the set of accessory genes varies between isolates. As expected, isolates from the same patients tend to share a similar set of accessory genes, which clearly distinguishes each patient’s isolates from the rest. Due to the clonal nature of the patient isolates, we selected a less redundant set of 32 reference isolates, which displays a similar distribution of sequence identity and shared gene content as the complete set (**Fig 1B, Metadata Summarized in Table 2**).

**Table 2.** Selected Metadata/MLST for Reference Isolates.

| patient ID | MSI | strain | Inflamed | Biopsy location | phylo group | MLST |
| --- | --- | --- | --- | --- | --- | --- |
| I | 0.98 | 4A | No | colon | D | 38 |
| I | 0.11 | 4C | No | colon | D | 38 |
| II-1 | 0.05 | 6J | No | colon | B1 | 224 |
| II-2 | 0.08 | 7C | No | colon | A | 216 |
| IV | 19.67 | 13B | Yes | terminal ileum | B2 | 141 |
| IV | 30.18 | 13N | Yes | terminal ileum | B2 | 141 |
| IV | 21.04 | 14C | Yes | terminal ileum | B2 | 141 |
| IV | 29.88 | 14D | Yes | terminal ileum | B2 | 141 |
| IV | 4.93 | 16B | No | cecum | B2 | 141 |
| IV | 26.19 | 16F | No | cecum | B2 | 141 |
| VII | 0.04 | 28A | No | colon | A | 10 |
| VIII | 2.84 | 29J | No | colon | A | 46 |
| VIII | 2.48 | 30B | No | colon | A | 46 |
| VIII | 1.58 | 30E | No | colon | A | 46* |
| VIII | 0.19 | 30G | No | colon | A | 46 |
| XI-1 | 0.22 | 42F | No | colon | B2 | 144 |
| XI-2 | 0.23 | 42J | No | colon | G | 117 |
| XI-2 | 1.53 | 44C | Yes | colon | G | 117 |
| XII | 1.68 | 45B | Yes | pouch | E | 11 |
| XII | 1.50 | 47H | Yes | pouch | E | 11 |
| XIII | 0.38 | 51C | No | colon | E | 11 |
| XX | 0.02 | 74D | Yes | cecum | A | 2223 |
| XX | 2.36 | 75P | No | colon | A | 2223 |
| XXIV | 0.10 | 90B | No | colon | D | 130 |
| XXXIII-1 | 0.21 | 125D | No | terminal ileum | B1 | 8492 |
| XXXIII-1 | 4.13 | 128H | No | colon | B1 | 8492 |
| XXXIII-2 | 0.13 | 126H | No | anastomosis | A | 607 |
| XXXIII-2 | 10.02 | 127E | No | colon | A | 607 |
| XLII | 1.56 | 161F | No | colon | B2 | 131 |
| XLII | 1.20 | 161J | No | colon | B2 | 131 |
| XLII | 2.72 | 164I | No | terminal ileum | B2 | 131 |
| XXXIX |  | 150G |  |  | B2 | 95 |

Despite the clonal tendency of isolates, different *E. coli* strains (according to MLST) existed in three of the patients with CD (Patient II, XI, and XXXIII), which has been reported for other CD patients (23). When multiple strains existed within a single patient, they belonged to alternate phylogroups. For example, isolates from patient XI belong to two clonal populations represented by phylogroup B2 (closest to the NRG 857c AIEC strain) and phylogroup G. The phylogroup G isolates from patient XI belong to the main sequence type (ST117) for the group. Previous epidemiologic studies on ST117 isolates suggested their lineage is associated with poultry, but they can cause extra-intestinal disease in humans (25). Similarly, two patients presented with isolates from both phylogroups A (ST216 in patient II and ST607 in patient XXXIII) and B1 (ST224 in patient II and ST8492 in patient XXXIII), with one of the B1 sequence types (ST224) reported as a high-risk lineage with antibiotic resistance found in healthy chickens (26).

### Antibiotic resistance compromises gentamicin protection assays for AIEC invasion

All AIEC isolates were initially characterized by field-standard gentamicin protection assays to measure their ability to invade epithelial cells (**Fig 2A**). The mean survival index for this assay (MSI_gent_) displayed a broad range of values (0.022 – 35) across all isolates (**Table S3**), with the highest invasion scores stemming from clonal isolates of a single patient (IV). To assess if any protein-coding genes segregate with bacterial invasion, we correlated the genetic differences between isolates with the MSI_gent_ scores (**Table S4**). Two of these correlations are shown in **Fig 2B**, with the top-ranked protein annotated as aminoglycoside N-acetyltransferase AAC(3)-VIa (AAC-VIa). This enzyme inactivates aminoglycoside antibiotics, including gentamicin, by acetylating the 3-amino group, providing gentamicin resistance in bacteria that express the protein (27). The identification of this co-segregating gentamicin resistance gene suggests the inference of invasion phenotype from the MSI_gent_ scores is compromised. However, we provide functional annotations for proteins encoded by all genes correlating with MSI_gent_ scores for completeness (**Table S5**).

**Figure 2.**
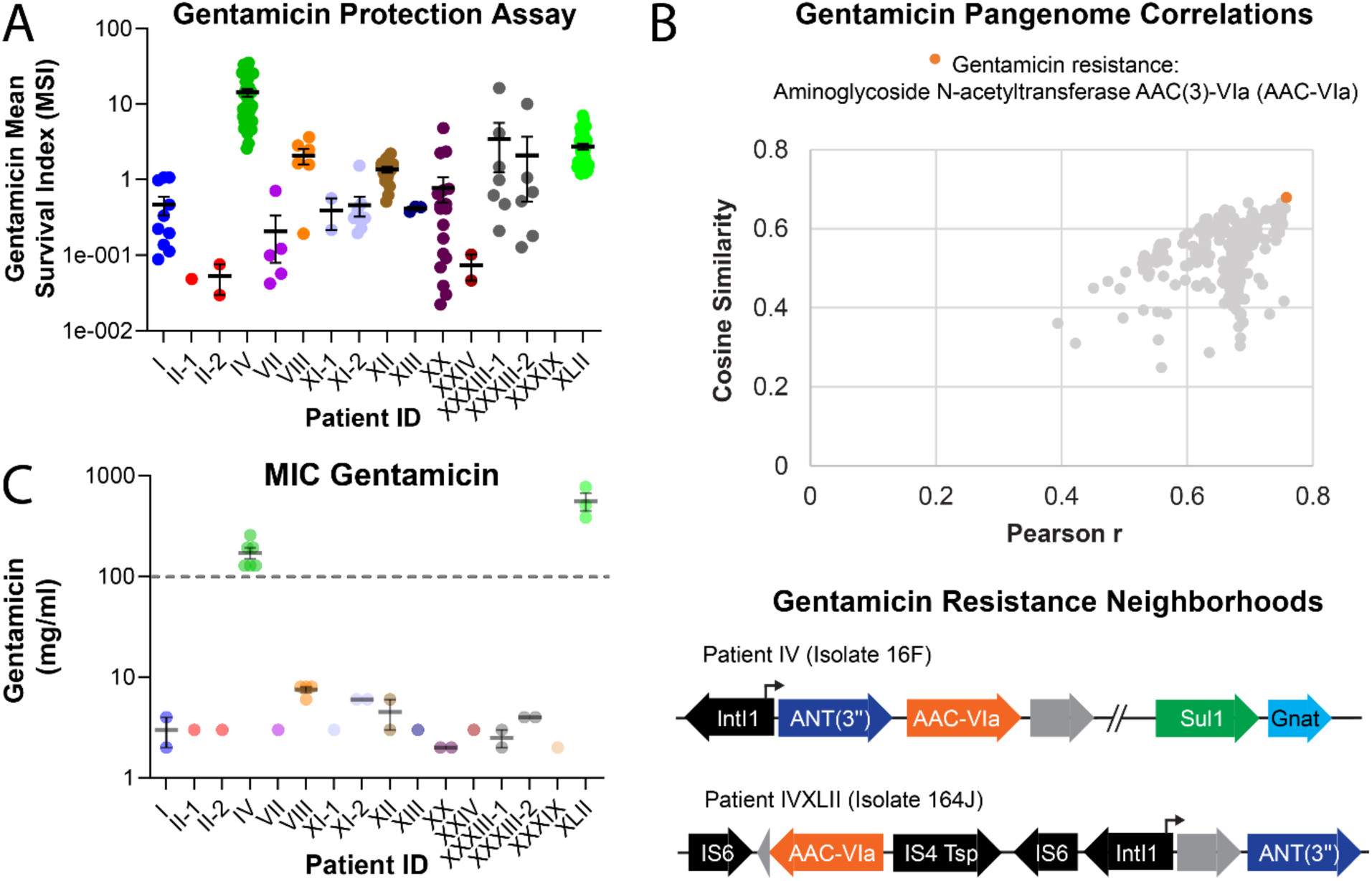
Acquisition of gentamicin resistance by *E. coli* isolates hinders invasion assay. **(A)** Mean survival index (MSI) measured by gentamicin protection assay is depicted for patient isolates (colored circles by patient, as in panel A of Figure 1). Isolates were grouped according to MLST type. The mean (horizontal line) and standard error of the mean are depicted for all grouped scores. **(B)** Pangenome distance correlation with LogMSI, as measured by cosine similarity (X-axis) is compared to that measured by Pearson correlation (Y-axis). Only significantly correlated proteins (by Pearson and QTL statistics; see methods) are shown. Proteins with similar functions are colored and labeled above the graph. The top-ranked protein is labeled with its genome neighborhood from patients IV and XVII, illustrated below. **(C)** Minimum Inhibitory Concentration (MIC) for gentamicin measured for representative patient isolates (colored circles, as in panel A). The dashed line indicates the concentration used in antibiotic protection assays.

The AAC-Vla gene is present in *E. coli* isolates from two patients (IV and XLII), with each located near a class 1 integron integrase (IntI1, **Fig 2B**). These integrons enable the horizontal spread of genetic determinants across clinical isolates and play an important role in the transmission of drug resistance (28, 29). The AAC-Vla neighborhood of patient IV represents a classic arrangement of the IntI1 adjacent to an array of gene cassettes, which include the gentamicin resistance gene as well as another aminoglycoside antibiotic that nucleotidylylates streptomycin at the position 3’’ hydroxyl group (ANT (3’’)), according to the Comprehensive Antibiotic Resistance Database (CARD) (30). CARD also helped us find another gene related to a known sulfonamide-resistant dihydropteroate synthase (Sul1) and an N-acetyltransferase (gnat) that may also modify an antibiotic in the upstream region to AAC-Vla. Thus, isolates from patient IV likely acquired gentamicin resistance through IntI1-mediated recombination into the integron array of gene cassettes. The AAC-VIa neighborhood of patient XLII, despite being near an IntI1 and its cassettes, is flanked by IS6 transposases (**Fig 2B**). Thus, the IS6 (or adjacent IS4) transposase could have mediated the acquisition of gentamicin resistance as a passenger gene in this isolate (31).

Given the identification of gentamicin and other potential antibiotic-resistance cassettes in the isolates of two patients, we comprehensively searched for known antibiotic-resistance genes encoded by all the sequenced genomes to predict each isolate’s drug resistance (**Table S6**). Isolates from all patients are predicted to be resistant to beta-lactam, and all but patient VII are likely resistant to Fosfomycin. In addition to their resistance to gentamicin, isolates from patient XLII include probable resistance to 10 of the 15 antibiotics considered. Given this propensity for antibiotic resistance in our isolates, we screened a subset of these strains with additional antibiotics to assess for their minimum inhibitory concentration (MIC) required to prevent growth. Consistent with the predicted gentamicin resistance, patients IV and XLII exhibit MICs for gentamicin higher than the concentrations used in the protection assays (**Fig 2C**). Even more, many other isolates exhibited intermediate resistance (>2mg/ml) to gentamicin.

### Development of an alternate antibiotic protection assay to estimate invasion

The expanded antibiotic screen confirmed resistance among the isolates to additional drugs (**Table 3**). Patient VIII isolates, which were predicted to be resistant to kanamycin by genomic sequences, were also resistant experimentally. Alternately, patient IV isolates were predicted to be resistant to ampicillin, but the bacteria exhibited low MIC when grown in the presence of the antibiotic. A similar difference between prediction and experimental measurement was seen with Patient XLII isolates, which were resistant to ampicillin by experimental results but not predictable by sequence. These discrepancies highlight the complicated nature of antibiotic resistance of bacteria and the necessity for experimental screening prior to choosing antibiotics in protection assays for invasiveness.

**Table 3.** Antibiotic resistance screen of AIEC isolates.

| Strain | Patient ID | Antibiotic Resistance Profile* |  |  |  |
| --- | --- | --- | --- | --- | --- |
|  |  | Gentamicin | Ceftazidime | Ampicillin | Kanamycin |
| 4A | I | I | S | S | S |
| 4C | I | S | S | S | S |
| 6J | II-1 | I | S | S | S |
| 7C | II-2 | I | S | S | S |
| 13B | IV | R | S | S | I |
| 13N | IV | R | S | S | S |
| 14C | IV | R | S | S | I |
| 14D | IV | R | S | S | I |
| 16B | IV | R | S | S | I |
| 16F | IV | R | S | S | I |
| 28A | VII | I | S | S | S |
| 29J | VIII | I | S | S | R |
| 30B | VIII | I | S | S | R |
| 30E | VIII | I | S | S | R |
| 30G | VIII | I | S | S | R |
| 42F | XI-1 | I | S | S | S |
| 42J | XI-2 | I | S | S | S |
| 44C | XI-2 | I | S | S | S |
| 45B | XII | I | S | S | S |
| 47H | XII | I | S | S | S |
| 51C | XIII | I | S | S | S |
| 74D | XX | R | S | R | R |
| 75I | XX | R | S | R | R |
| 90B | XXIV | R | S | R | R |
| 125D | XXXIII-1 | S | S | S | S |
| 128H | XXXIII-1 | S | S | S | S |
| 126H | XXXIII-2 | I | S | S | S |
| 127E | XXXIII-2 | I | S | S | S |
| 161F | XLII | I | S | S | S |
| 161J | XLII | S | S | S | S |
| 164I | XLII | I | S | S | S |
| 150G | XXXIX | S | S | S | S |

\*MIC ≥ susceptible threshold and ≤ resistance threshold; S = susceptible, R = resistant, and I = intermediate

Using the expanded antibiotic screen, we initially tested alternative antibiotics for their ability to penetrate the host cell, as this would compromise our invasion assays. Using multiple classes of antibiotics, including ceftazidime, cefepime, ertapenem, and amikacin, we found that only amikacin did not penetrate Caco-2 cells at 10x MIC (**Supplemental Figure S1**). We also observed a lack of resistance to amikacin among our isolates and a lack of toxicity for Caco-2 (**Supplemental Figure S2**). We therefore developed a protection assay using this antibiotic with representative strains defined by the phylogenetic tree and MLST. Interestingly, the patient isolates with acquired gentamicin resistance were on antibiotic treatments at the time of their colonoscopy (**Table S7**). Representative isolates from six patients (XI-1, XI-2, XII, XIII, XX, and XXIV) and the positive control, NRG857c, were significantly different from the negative controls in this amikacin protection assay (**Fig 3A**, **Table S8**), suggesting these strains are invasive. Amikacin protection measured for isolates from another patient (II-1) were borderline (0.035 Adjusted P-value, noted by a single *), which we conservatively classified as non-invasive. An isolate from an asymptomatic control patient (XXXIX) with a normal colonoscopy and without evidence of inflammation or colonic polyps was not significantly different from the negative controls for invasion (**Fig 3A**).

**Figure 3.**
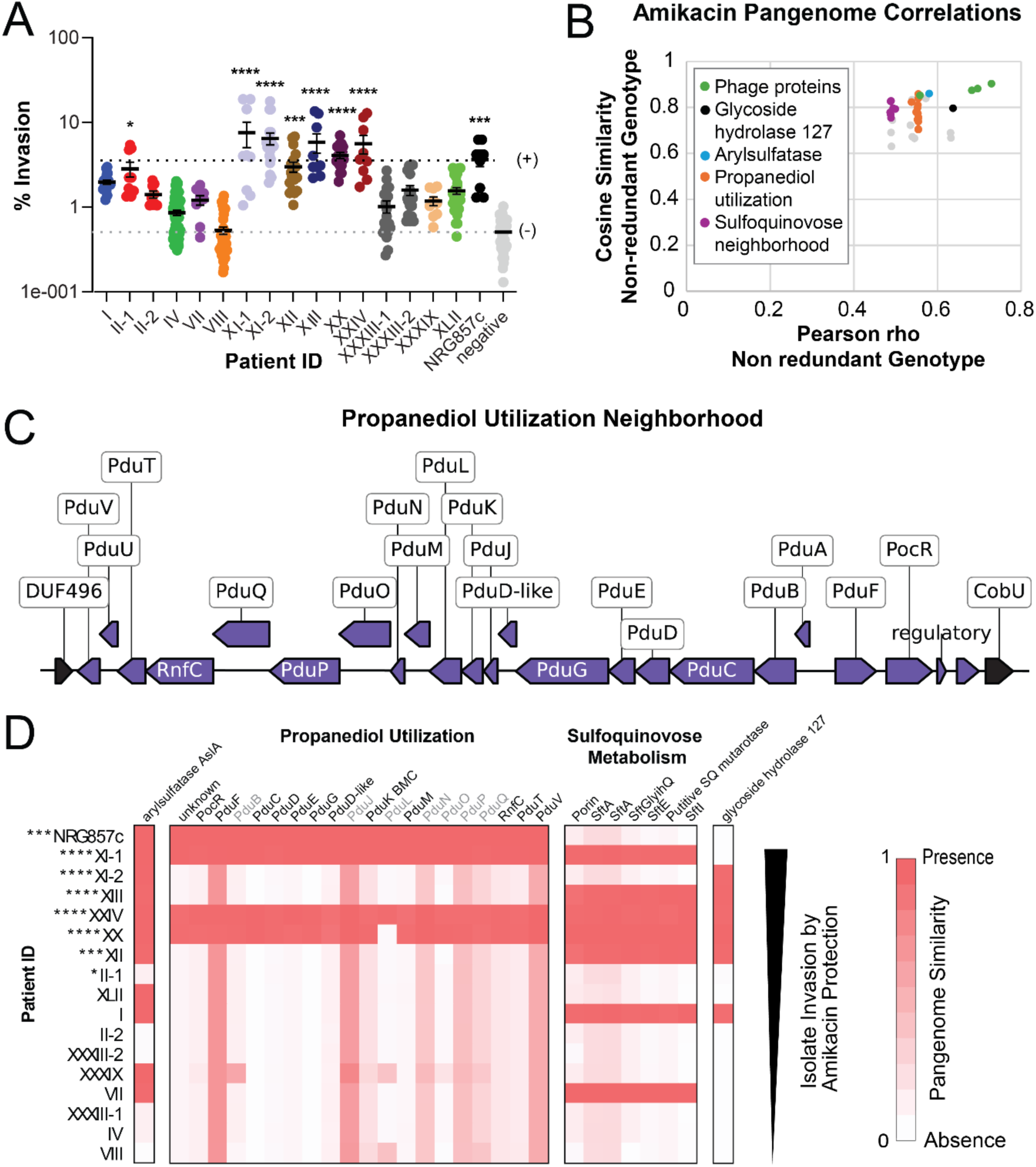
Amikacin protection assay for isolate invasion. **(A)** Amikacin invasion assay for reference isolates (colored circles, as in Figure 2) with mean (horizontal line) and standard error of the mean depicted for MLST grouped reference isolates. Horizontal dashed lines represent the means for the NRG 857c positive control (black) and all negative controls (gray). Patients with percent invasion scores that are significantly different from the negative controls are indicated by (*). **(B)** Pangenome distance correlation with control scaled amikacin protection scores are measured by cosine similarity calculated for non-redundant isolates (X-axis) and Pearson correlation calculated for non-redundant isolates (Y-axis). Only significantly correlated proteins (by Pearson and QTL statistics; see methods) are shown. Proteins with similar functions are colored and labeled in the graph insert. **(C)** The propanediol utilization gene neighborhood (purple arrows labeled above) is depicted for the NRG857c AIEC reference strain, with the flanking genes indicated by black arrows. **(D)** Pangenome similarity of patient isolates to AslA, propanediol utilization gene neighborhood, SQ metabolism neighborhood, and Gly127; with scaled scores between 0 (absent) and 1 (present). Patient isolates are ordered below the NRG857c AIEC positive strain from highest (top) to lowest (bottom) amikacin protection scores.

### Co-segregation of virulence factors with invasion phenotype

We sought to identify genes that correlate with one of the phenotypes used to define AIEC strains, namely their invasiveness (MSI_amk_) as measured by the amikacin protection assays. These gene correlations (**Table S9**) were less significant than those for gentamicin (likely due to the lowered number of isolate genomes), and the significant gene set included fewer proteins (**Table S10** and **Fig 3B**). Among these, the most significantly correlated proteins are unknown phage proteins, a GH127 family protein, and AslA. Two sets of proteins functioning in propanediol utilization and SQ metabolism were also correlated to the invasion of Caco2 cells. The propanediol utilization proteins belong to a genome neighborhood (**Fig 3C**) present in the positive control strain for invasiveness, i.e., NRG857c, which is commonly found in ileal lesions of CD patients. This neighborhood was noted as unique to the NRG857c strain by comparative genomic analysis to other pathogenic and commensal *E.coli* (32). Most propanediol utilization proteins are identified as correlated with the invasion phenotype, as they belong to the same genomic neighborhood; exceptions (e.g., PduQ and PduJ) exhibit intermediate similarity to proteins participating in ethanolamine utilization, representing a paralogous metabolic pathway to propanediol utilization that is present as a core component of all *E. coli* isolates.

The propanediol utilization neighborhood was found in invasive isolates from three of six patients tested (patients XI-1, XX, and XXIV), and these isolates displayed significantly higher invasion than the negative control (**Fig 3B & C**, significance marked by *). The presence of this genomic island in patient isolates with different phylotypes (**Fig 1A**, phylotypes B2, A, and D) highlights its mobile nature and suggests its acquisition contributes to the invasion phenotype. Isolates from three other patients that were observed to invade Caco-2 cells in the amikacin protection assay lacked this propanediol utilization neighborhood (patients XI-2, XII, and XIII). Two of these patients encode SQ metabolism genes, which also significantly correlate with the invasion phenotype. Finally, all our sequenced invasive isolates have acquired AslA, and all but patient XI-1 and the positive control NRG 857C encode GH127. In sum, different isolates seem to have acquired alternative invasion strategies, and they collectively enable these isolates to colonize human epithelial cells and ultimately cause disease.

## Discussion

This study analyses bacterial isolates from IBD patients using comparative genomics to identify factors that can contribute to cell invasion. We utilized Illumina short reads and Nanopore long reads to obtain high-quality genome assemblies for our isolate collection. Consistent with previous studies, we show that there is clonality of isolates within each patient regardless of biopsy location, apart from three IBD patients that had more than one dominant isolate. Since bacterial invasion of enteric cells can be a key feature of virulent strains of *E.coli* in IBD, we further characterized the invasion capacity of the sequenced strains using the established gentamicin protection assay. Surprisingly, several of our strains carried genetic features of gentamicin resistance. Therefore, we developed an alternative antibiotic protection assay, based on sensitivity to amikacin. Through this approach, we were able to identify isolates that survived exposure to amikacin. By correlating each strain’s amikacin protection assay score (a proxy for the invasiveness of each isolate) with their genomic sequence, we identified genes involved in propanediol utilization proteins and sulfur metabolism that might be novel *E. coli* virulence factors required for invasion.

The observation that a sulfur metabolizing pathway (SQ metabolism) and enzyme (AslA) are among the top co-segregating proteins with the invasion phenotype of *E. coli* isolates suggests colonic sulfur metabolism might contribute to invasion and lead to inflammation associated with IBD. While the extent to which the products of the SQ pathway and the AslA reaction lead to H_2_S production by colonic microbiota remains undetermined, the gut microbiome seems to play a significant role in IBD pathology (33). The relationship between H_2_S and colonic health remains debated (34, 35). However, elevated H_2_S can exhibit toxic properties to colonocytes, induce inflammation, and increase intestinal permeability (34, 36). Sulfur is provided to the human body in the diet, mainly through protein consumption. It is conceivable that exogenous sulfate levels in the gut may contribute to metabolism-linked AIEC in some genetic backgrounds.

The longest stretch of syntenic co-segregating genes that we identified in our invasive *E. coli* isolates encoded propanediol utilization islands. Previous studies have also identified the propanediol utilization proteins as being enriched in pathogenic AIEC compared to control strains (32), and the pathway appears to contribute to AIEC invasiveness and inflammation (37, 38). Propanediol utilization occurs in a bacterial microcompartment (BMC) formed by a proteinaceous shell that sequesters the involved chemical reactions into a primitive organelle (39). The propanediol BMC concentrates low levels of the metabolic enzymes, as well as volatile reaction intermediates, to enhance pathway flux from fucose and rhamnose precursors through 1,2 propanediol and keep the levels of toxic propionaldehyde low (40). Such BMCs also allow enteric pathogens like *Salmonella* to colonize the mammalian gastrointestinal tract and outcompete native microbiota (41, 42). The presence of propanediol utilization islands in *E. coli* isolates from three of our patients suggests these bacteria also might take advantage of this niche carbon source for successful colonization and invasion during inflammatory colitis.

The co-segregation of SQ degradation islands in some patient isolates lacking propanediol utilization suggests alternate genetic strategies might contribute to invasion phenotype. SQ is a sulfonated monosaccharide in green vegetables that serves as a nutrient source for select microbes in the gut and can lead to increased microbiota-generated H_2_S levels (43). Bacteria that typically degrade SQ from green vegetables in the human gut are generally associated with a positive impact on human health, bringing into question the pathobiology of SQ metabolism in AIEC invasion. However, microbiome released H_2_S from the products of SQ catabolism can have detrimental consequences on colonic health, including opening the mucus barrier in the colon and allowing bacteria access to the epithelium lining (36). Another sulfur-metabolizing protein, AslA, co-segregates with invasion. While AslA has not yet been defined as a contributing factor to AIEC colonic invasion, it is considered an invasion factor for *E. coli* in brain microvascular endothelial cells (44). Sulfatase activity like that potentially catalyzed by AslA can provide access for the bacteria to otherwise heavily sulfated colonic mucin glycans (45). This activity might provide invading bacteria with glycans as a nutrient source, with access to the intestinal epithelium, or may increase sulfate levels for colonic sulfur metabolism. Another more universally encoded enzyme among invasive isolates, GH127, breaks down l-arabinofuranose-β1,2-l-arabinofuranose as a potential nutrient source, perhaps also acquired from plant polymers in the diet (46).

Comparison of gene content between different isolates in this study also identified a considerably wider range of shared orthologs compared to sequence identity, with as little as 74% of shared genes among our isolates compared to 96-99% average nucleotide identity. Genetic variability can result from horizontal genetic transfer, genetic diversification, and gene loss, especially under the pressure of antibiotic selection. IBD patients often have high antibiotic exposure and thus may be at risk for acquiring antibiotic-resistant organisms. Antibiotics such as ciprofloxacin and metronidazole have been used to treat IBD, and the prevalence of patients that have been exposed to these antibiotics is as high as 15% (47–49). Further, IBD patients have a higher incidence of conditions that require antibiotics, including bacterial overgrowth manifesting from gut microbial dysbiosis and an increased prevalence of *Clostridium difficile* infections (49, 50), abscesses, and wound infections (51, 52). Biologics and immunomodulators, the current mainstay of CD therapy, have added additional exposure to antibiotics, as they predispose patients to infection.

Gentamicin protection assays are the current standard of practice to test for invasion of bacterial isolates and phenotypically define them as AIEC (17, 18). However, in this study we found that 28.1% of the clinical isolates survived gentamicin protection. Using whole genome sequencing of this data set, we then searched for genes that correlate with the results of the gentamicin invasion assays and found AAC-Vla, an enzyme that provides resistance to gentamicin, to be the top-ranked gene. Based on these results, we concluded that it was likely that the survival of these strains in the presence of gentamicin was not due to invasion, but to gentamicin resistance. With additional testing, we found resistance to multiple antibiotics among our clinical isolates, highlighting the necessity of determining MIC for a broad panel of antibiotics before performing cell protection assays as a proxy for invasion. This practice will contribute to eliminating false positives for the identification of invasive phenotype for AIECs.

Taken together, these findings highlight the information gained through sequencing. Despite the ever-increasing availability of whole genome sequences and the power of comparative genomics, genetic markers for the AIEC pathotype are lacking. Only a few virulence factors have been described in AIEC, including adhesins such as fimbriae (53), siderophore-associated uptake proteins (37), and capsule synthesis proteins (54). However, none of these are specific to the pathotype. Therefore, our approach using both long read and short read sequencing paired with accurate phenotypic characterization was able to identify putative virulence genes in strains of *E. coli* from IBD patients.

## Methods

### Collection of bacterial isolates and Measurement of Mean Invasion Index

This study involved the prospective collection of clinical *E. coli* isolates from eligible subjects undergoing colonoscopy, ileoscopy, or flexible sigmoidoscopy at a single center as described (21). Briefly, all participants signed informed consent, and the Emory Institutional Review Board approved this study. Patients with clinically confirmed IBD were recruited between July 2002 and August 2005, and four biopsies were obtained from each patient during outpatient colonoscopy. As previously described, As previously described, biopsies were incubated in Dulbecco’s Modified Eagle’s Medium (DMEM, Invitrogen, Carlsbad, CA, USA) supplemented with 100 μg/ml gentamicin (Invitrogen) for 1 hour, washed three times in PBS and lysed in 1% Triton-X-100/PBS. Aliquots were cultured on MacConkey agar plates at 37°C overnight and lactose-fermenting colonies were enumerated and propagated in LB broth for 4 h at 37°C under aerobic conditions. Individual clones were stored in 50% glycerol at −80°C until further use.

### Isolate sequencing

Total DNA was extracted from overnight cultures of clinical isolates (in Luria-Bertani broth at 37 degrees Celsius) using the Qiagen DNeasy Blood & Tissue kit. We prepared a paired-end library for Illumina sequencing and a long-DNA library for Nanopore sequencing for each isolate. After sheering the gDNA with NEBNext Ultra II FS kit, we prepared paired-end libraries with the NEBNext® Ultra™ DNA Library Prep kit for Illumina®. We pooled libraries of different samples and sequenced them using the Hiseq X ten sequencing service from Azenta, targeting a 100-fold coverage (0.5Gb) per isolate. For long-read libraries, we used the LSK109 kit with Native Barcodes from kits EXP-NBD114, EXP-NBD114, and EXP-NBD196 for multiplexing. We made the following modifications to the standard protocol for the LSK109 kit: (1) we extended the incubation time for DNA repair and end-prep to 1 hour at 20°C; (2) we determined the needed Adapter Mix II (AMII) volume based on the amount of pooled DNA during the adapter ligation step and used 6.0 µL AMII per 1.0 µg DNA. We sequenced the pooled libraries using the MinION flow cell R9.4.1, targeting 100-fold coverage (0.5Gb) per isolate.

### Genome assembly and annotation

Illumina sequences were processed with Trimmomatic 0.38 (55) to remove sequencing adapters, and low-quality bases (quality score < 20) and discard reads shorter than 30bp after such trimming. We performed base-calling, demultiplexing, and adaptor removal with Guppy, a software provided by Nanopore. Combining the Illumina and Nanopore reads, we assembled the genome of each isolate using Unicycler (56) with default settings. We used the NCBI Prokaryotic Genome Annotation Pipeline (57) with default settings to assign taxonomy to each isolate, predict protein-coding sequences in the genomes, and annotate each protein with function. Additionally, we employed the Artificial Intelligence tools ProteInfer (58) and ProtNLM (59) to obtain additional function annotations.

### Isolate phylogenetics, MLST assignment, and genetic diversity estimation

The assembled genomic sequences from each isolate were submitted to the M1CR0B1AL1Z3R web server (60) to extract orthologous sets of putative open reading frames and reconstruct a phylogenetic tree from the common set of orthologs. The phylogenetic tree was visualized using the interactive tree of life (iTOL) v5 online tool (61). Multilocus sequence typing (MLST) analysis was carried out by searching (62) our genomes against protein sequences from the *Escherichia* sequence typing database at PubMLST (63) and comparing results to known *E. coli* (Achtman) allelic profiles. To determine the phylogroup, genomic sequences of representative isolates from each MLST group were submitted to ClermonTyping (25, 64). To estimate the genetic diversity among isolates, we used fastANI (65) to estimate the average nucleotide identity and fraction of shared genes between each pair of isolates.

### Building a Pangenome Proteome Reference

We built a pangenome reference protein set to enable comparative analysis. We classified proteins in all the *E. coli* isolates by OrthoFinder (v2.5.4) (66). We used MAFFT (67) to align proteins in each orthologous group and identified positions shared by 25% of proteins. Proteins covering at least 95% of such shared positions were considered candidate reference proteins. In rare cases where no protein covered 95% or more of the shared positions, we considered proteins covering at least *C*_*max*_ − 10 shared positions as candidates, where *C*_*max*_ is the largest number of shared positions covered by a protein. Among these candidates, we selected one reference per orthologous group, prioritizing proteins from the reference strain NRG857c or isolates with higher invasiveness as determined by gentamicin protection assays. We calculated the number of other proteins between the coding genes of every protein pair in every *E. coli* genome, i.e., *dist*_*x*,*y*_. We computed the genomic distance between each pair of reference proteins *X* and *Y* as *DIST_X_*_,*Y*_ = *median* (*dist*_*x*,*y*_), where *x* and *y* are proteins from the same species and belong to the orthologous groups represented by *X* and *Y*, respectively. The high dimensional matrix with *DIST_X_*_,*Y*_ for all pairs of reference proteins was reduced to a one-dimensional vector using the manifold.MDS function from the sklearn module of Python; values in this vector were used to order proteins in the pangenome reference in **Fig. 1**.

### Pangenome Protein Distances and Correlation with Invasion

We searched against the proteome of each isolate with the pangenome reference proteins as queries using BLAST (68) (e-value cutoff: 1000) and extracted the bitscore from the top hit to each isolate. For each pangenome reference against proteome, we calculated a similarity score based on the bitscore of the top hit (*P*_*bs*_) and the bitscore of aligning the query against itself (*M*_*bs*_) as *genoS* = *P*_*bs*_/*M*_*bs*_. The *genoS* scores of each reference protein over select isolates were then compared to their phenotypic scores (*phenoS*) using cosine similarity, Person’s correlation, and Spearman’s rank correlation implemented in Python Scipy module. In addition, we performed Quantitative Trait Locus (QTL) analyses to identify genes whose presence or absence in an isolate can predict its phenotype. For each protein, we partitioned the isolates into two groups based on whether it is present in the proteome; the correlation between this gene’s presence/absence (genotype) with phenotype is evaluated by student’s T-tests on *phenoS* of the two groups defined by the genotypes.

For gentamicin protection assays, we transformed the mean survival index (MSI_gent_) to a log scale to obtain *phenoS*. For the alternative invasion assay, the pre-scaled percent invasion scores (MSI_amk_, see below) were used as *phenoS*. Because such correlations are only meaningful if *genoS* and *phenoS* for different isolates have diverse values, and we constrained our calculations to proteins satisfying three criteria: (1) *genoS* shows at least 3 different values; (2) max(*genoS*) − min(*genoS*) > 0.01 · *mean*(*S*); (3) max(*phenoS*) − min(*phenoS*) > 0.01 · *mean*(*phenoS*). We were also concerned that isolates from the same patient cannot be treated as independent samples. Thus, in addition to analyzing the entire set of isolates with experimental data, we also assigned the isolates into groups based on phylogeny (**Tables S4 and S7**), averaged both *distS* and *phenoS* scores within each group, and repeated these analyses on the average values. To find proteins showing significant correlation with the phenotypes, we required Pearson and QTL Q-values (false discovery rate (69)) < 0.05 for correlations using all isolates and the Pearson and QTL P-value < 0.05 for correlations using averages of phylogenetic groups of isolates.

### Antibiotic resistance screening and development of alternate invasion assay

We used AMRFinderPlus (70) developed by NCBI to detect antibiotic resistance in the isolates using both proteins (-p) and genomes (-g), specifying *Escherichia* (-O) as the organism. MICs of antibiotics were determined using MIC Test Strips (Liofilchem). To perform antibiotic protection assays, Caco-2 cells were seeded on collagen-coated plates. To minimize basolateral infection, cells were allowed to differentiate for 3 days before infection. Each monolayer was infected in triplicate at a multiplicity of infection (MOI) of 10 bacteria per epithelial cell (71). After a 3-hour incubation period at 37 °C, infected monolayers were washed with PBS to remove non-adherent bacteria. Extracellular bacteria were eliminated by one-hour incubation with DMEM containing Amikacin at 10x MIC of the isolates. Monolayers were washed, and monolayer integrity was confirmed microscopically after the final wash. Epithelial cells were lysed in 1% Triton-X 100 in PBS (Sigma). Samples were then serially diluted and plated on LB agar plates to determine the number of bacterial colony-forming units (CFU) recovered from the lysed monolayers. Significant epithelial invasion was determined by comparisons to pathogenic *E. coli* strain NRG 857c (positive control) and nonpathogenic strains DH5α, MP7, and MG1655 (negative controls).

Amikacin invasion scores were performed in triplicate, reporting the CFU/ml and percent invasion score (PIS) for each reference isolate (**Supplemental Figure S3**). PIS was defined as the percentage of intracellular bacteria at 1 hour after amikacin treatment relative to that of the original inoculum. Invasion assays were repeated three times for each isolate, and the averages were plotted as a group (Prism 10.1.1) according to phylogeny. Amikacin PISs were scaled according to the controls using two methods. The first method (pre-mean scaled score) applies such scaling prior to averaging. The PIS for each assay (*PIS_X_*) was scaled according to the following equation: (*PIS_X_* − *PIS*_*ne*g_)/(*PIS*_*pos*_ − *PIS*_*ne*g_), where *PIS*_*pos*_ is the PIS for the positive control (NRG 857c) and *PIS*_*ne*g_is the average PIS for the four negative controls (DH5α, MP7, MP13 and MG1655). The three scaled scores for each experiment were then averaged, reporting the pre-scaled mean for each reference isolate. For the second method (post-mean scaled score), outliers were removed from PISs grouped by patient and phylogeny using the built-in ROUT method (Q = 1%, 5 outliers) of GraphPad. The mean PIS across replicates after outlier removal was calculated for each isolate, followed by rescaling by the same equation as the first method. Scores from both methods correlate (**Supplemental Figure S2C**), with a Pearson’s correlation coefficient of 0.97. Both pre-mean and post-mean scaled scores identified similar co-segregating genes. We chose to illustrate the pre-mean scaled scores (denoted as MSI_amk_).

We tested the significance of invasion phenotype by ordinary one-way ANOVA in GraphPad, comparing all PISs grouped by patient and phylogeny (without removing outliers). Each group was compared to the mean of the negative controls using Dunnett’s multiple comparison test, and P-values less than 0.05 were reported.

## Abbreviations

AIEC: adherent-invasive *Escherichia coli*
IBD: inflammatory bowel disease
*E. coli*: *Escherichia coli*
CD: Crohn’s Disease
SQ: sulfoquinovose
AslA: arylsulfatase
GH127: glycoside hydrolase family 127
USCGs: universal single-copy genes
HGT: horizontal gene transfer
OG: orthologous group
MSI: mean survival index
AAC-VIa: N-acetyltransferase AAC(3)-VIa
IntI1: Class1 integron integrase
CARD: Comprehensive Antibiotic Resistance Database
Sul1: sulfonamide-resistant dihydropteroate synthase
gnat: N-acetyltransferase
MIC: minimum inhibitory concentration
BMC: bacterial microcompartment
QTL: Quantitative Trait Locus
MOI: multiplicity of infection
CFU: colony forming units
PIS: percent invasion score

## Notes

### Competing Interest Statement

The authors have declared no competing interest.

