## Supplemental Figures for "Genetic and microbial analysis of invasiveness for *Escherichia coli* strains associated with inflammatory bowel disease"

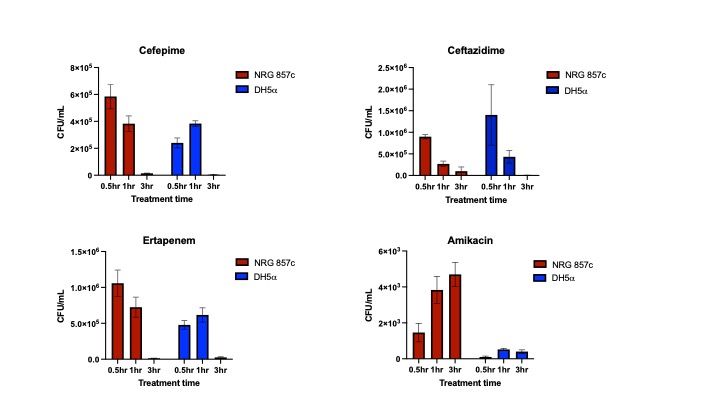

**Supplemental Figure S1. Ability of antibiotics to penetrate Caco-2 cells.** Antibiotic protection assays were performed by adding 10x minimal inhibitory concentration (MIC) of each antibiotic to Caco-2 cells which were pre-treated with bacteria (NRG857c vs DH5α). Colony forming units (CFU)/mL represents intracellular bacteria (y-axis) over antibiotic treatment time (x-axis).

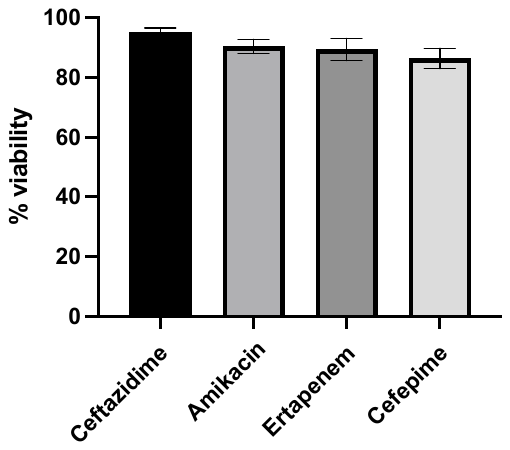

**Supplemental Figure S2. Cell viability of Caco-2 cells after antibiotic exposure.** Differentiated Caco-2 cells were incubated in media with 10x minimal inhibitory concentration of each antibiotic (ceftazidime, amikacin, ertapenem, and cefepime) for 7 hours. Cell viability was then measured using trypan blue.

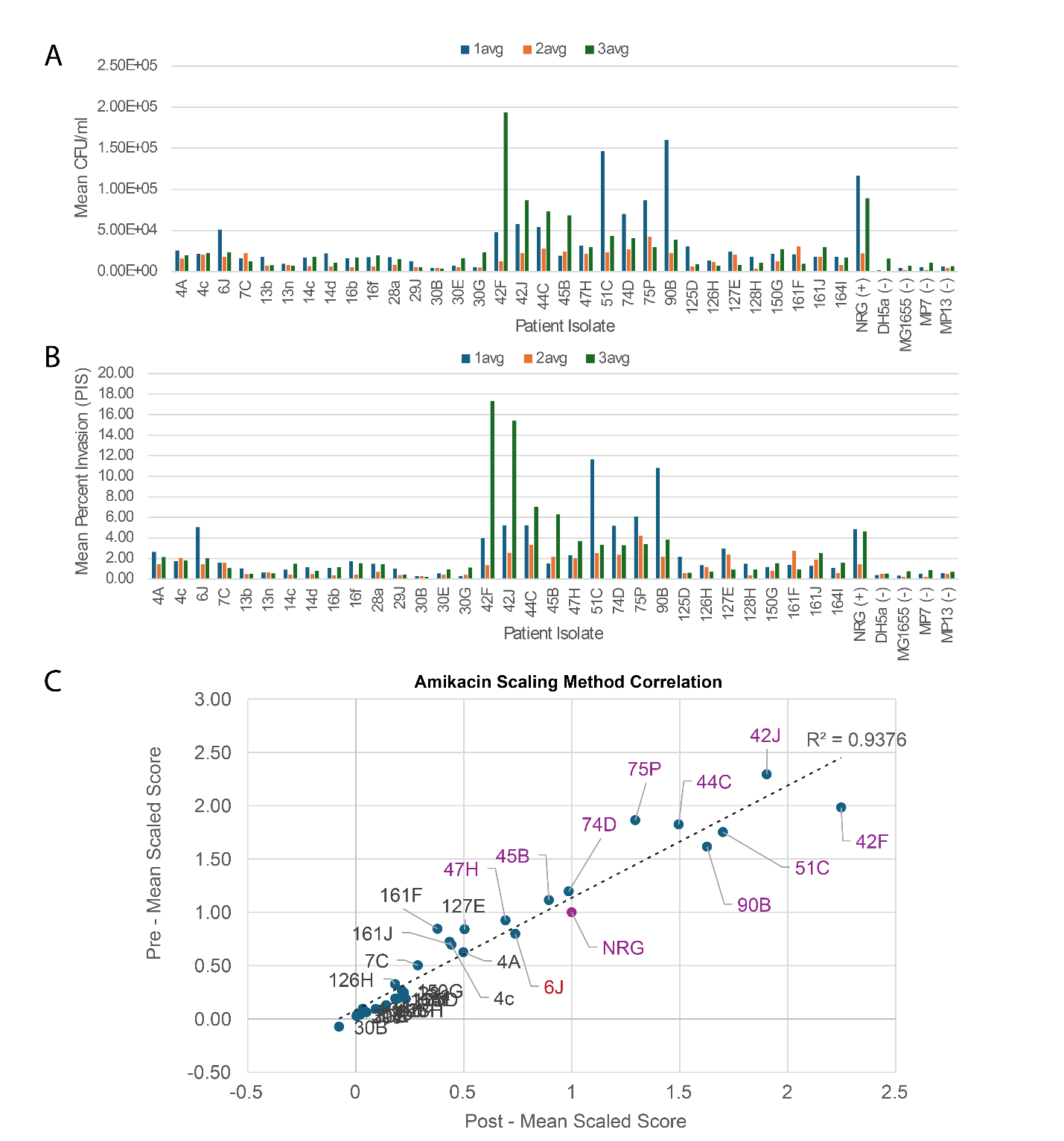

**Supplemental Figure S3. Amikacin protection assay scoring and transformation.** **A)** Calculated CFU/ml for indicated patient isolates and negative controls (-) and **B)** mean percent invasion (PIS) of Caco-2 cells measured as percentage of intracellular bacteria at 1 hour after amikacin treatment relative to that of the original inoculum. Each bar represents the average of three PIS. The colors indicate independent replicates (1Avg, 2Avg, and 3Avg). **C)** Correlation of PIS transformed using two methods: Pre-Mean scaling and Post-Mean scaling (see methods).
